# quickBAM: a parallelized BAM file access API for high throughput sequence analysis informatics

**DOI:** 10.1101/2021.10.05.463280

**Authors:** T. Anders Pitman, Xiaomeng Huang, Gabor T. Marth, Yi Qiao

## Abstract

**Motivation:** In time-critical clinical settings, such as precision medicine, genomic data needs to be processed as fast as possible to arrive at data-informed treatment decisions in a timely fashion. While sequencing throughput has dramatically increased over the past decade, bioinformatics analysis throughput has not, and consequently has now turned into the primary bottleneck. Modern computational hardware are capable of much higher performance than current genomic informatics algorithms can typically utilize, therefore presenting opportunities for significant improvement of performance. Accessing the raw sequencing data from BAM files, for example, is a necessary and time-consuming step in nearly all sequence analysis tools, however existing programming libraries for BAM access do not take full advantage of the parallel input/output capabilities of storage devices.

**Results:** In an effort to stimulate the development of a new generation of faster sequence analysis tools, We developed quickBAM, a software library to accelerate sequencing data access by exploiting the parallelism in commodity storage hardware currently widely available. We demonstrate that analysis software ported to quickBAM consistently outperforms their current versions, in some cases finishing an analysis in under 4 minutes while the original version took 1.5 hours, using the same storage solution.

**Availability and Implementation:** Open source and freely available at https://gitlab.com/yiq/quickbam/, we envision that quickBAM will enable a new generation of high performance informatics tools, either directly boosting their performance if they are currently dataaccess bottlenecked, or allow data-access to keep up with further optimizations in algorithms and compute techniques.

## INTRODUCTION

High throughput, genome wide next generation sequencing (NGS) have revolutionized precision medicine. As an example, NGS has now been implemented as a routine diagnostic modality in many pediatric subspecialty clinics for critically ill children admitted into the neonatal intensive care unit or pediatric intensive care unit(Elliott *et al*., 2019; Petrikin *et al*., 2015). And increasingly, genomics-guided precision medicine is helping advanced cancer patients who have exhausted standard-of-care options(Schwartzberg *et al*., 2017). In these settings, the amount of data analyzed is small compared to large cohort studies, involving usually one to a few tumor samples and a paired normal sample from the same patient. However, fast analysis turnaround is of critical importance. Furthermore, after the optimal treatment is identified, it still takes significant time to coordinate treatment access due to e.g. drug acquisition, compassionate care approval, clinical trial enrollment, or insurance authorization. It is therefore significant that the informatics analysis tasks, which have surpassed sequencing as the primary bottleneck, are to be as fast as current computer hardware can make possible.

The BAM file format(Li *et al*., 2009) is the current *de facto* standard for storing sequencing data generated from NGS experiments. BAM files are the most common starting place for various downstream analyses. The BAM format is the compressed, binary version of the SAM format, which we designed as part of the 1000 Genomes Project(1000 Genomes Project Consortium *et al*., 2015) to reconcile the once many different formats of storing sequencing data. Subsequently, software libraries are created to provide APIs to access the sequencing reads contained in a BAM file. HTSLIB(Bonfield *et al*., 2021) is the file access layer from Samtools(Li *et al*., 2009), the software developed to perform many SAM / BAM file related operations. The main focus of HTSLIB is to provide high level abstractions so that the programming interfaces stay the same regardless of the underlying file format (be it SAM, BAM, or CRAM) or storage and transport media (local files, HTTP URLs, or cloud storage). BamTools(Barnett *et al*., 2011) and SeqLib(Wala and Beroukhim, 2017) focus on modern C++ API designs for ease of programming. While BamTools implements its own BAM parsing logic, SeqLib integrates HTSLIB as the access layer. These libraries perform file access in a single-threaded manner (HTSLIB does support multi-threaded decompression, but not file reading), which is a reasonable design choice when 1) informatics analysis is compute bottlenecked, and therefore cannot benefit from faster data access; or 2) the file storage and transport media are incapable of high performance, e.g. when the files are served over low-bandwidth network attached storages. However, these tools cannot take advantage of storage technologies that are capable of much higher levels of file I/O parallelization and data bus bandwidth.

Specifically, we are concerned with two generally available storage technologies. On premise, the Lustre distributed file system is capable of achieving very high aggregated bandwidth by striping files onto different computer nodes and hard drives. And on the cloud such as the commercial Amazon Web Services (AWS), it is already commonplace to instantiate nonvolatile memory express solid state drives (NVMe SSDs) as the primary storage media. These two types of storage solutions cover the majority of high performance computing facilities, and both provide enough parallelism to support much higher analysis throughput than currently utilized.

We propose a new approach for parallelized BAM file access (**Figure 1**). We developed quickBAM, which uses two strategies to parallelize data reading. First, when the bam file index (BAI) is available, we utilize the “fixed-bin” indices which contain the starting file offset of each 16-kb genomic window. Second, when the BAI is not available (unsorted / unindexed BAMs or the unmapped region in indexed BAMs), we use a heuristic scanner (see ***methods***) to directly locate multiple starting locations for parallel parsing. Since the majority of sequence analysis tasks (e.g. quality control, various types of mutation calling) involve reading BAM files, quickBAM has the potential to significantly shorten end-to-end analysis turnaround. quickBAM is freely available at https://gitlab.com/yiq/quickbam/ with extensive accompanying documentation available at https://yiq.gitlab.io/quickbam/.

**Figure 1.**
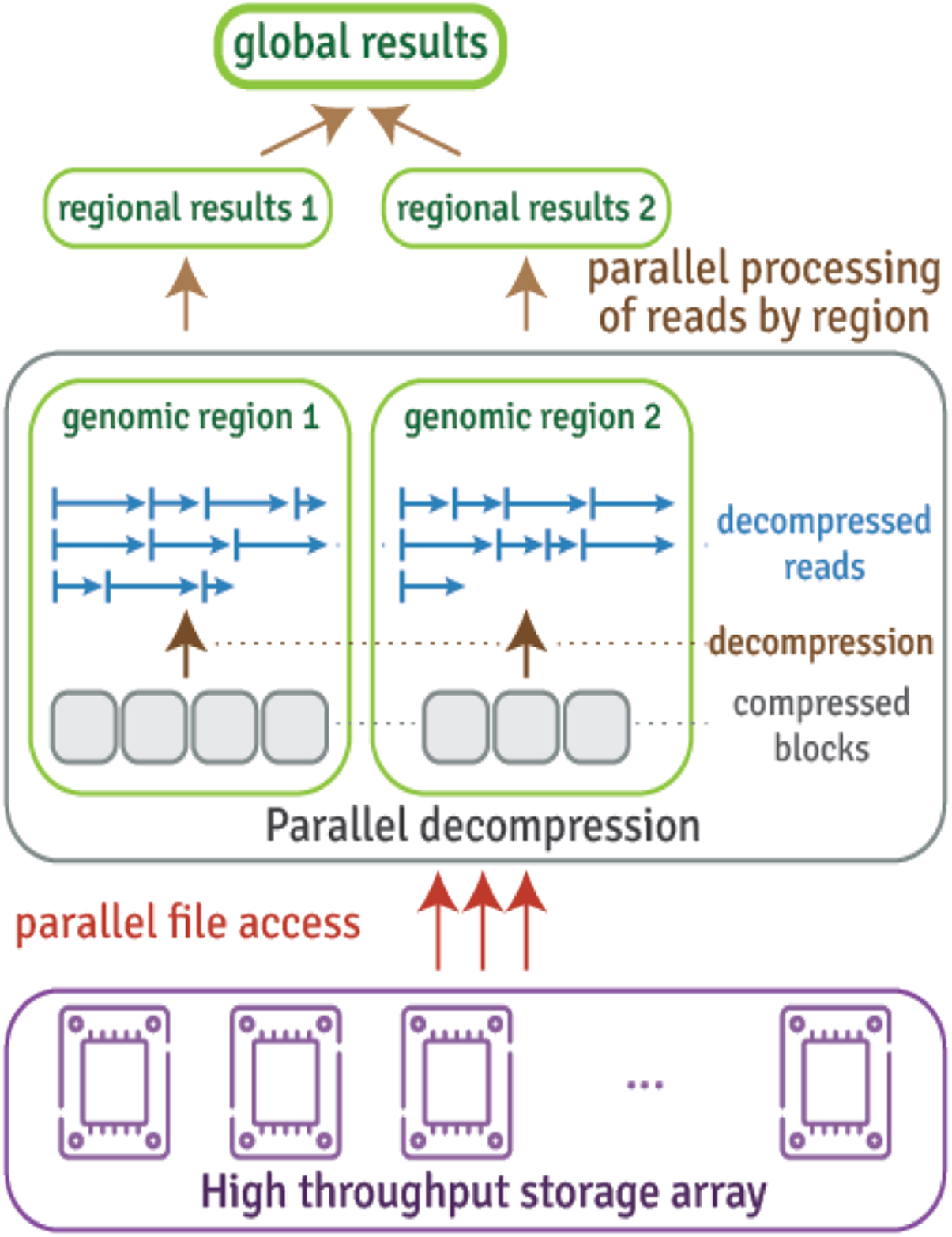
Parallel processing architecture using quickBAM. quickBAM utilizes the scatter / gather paradigm to parallelize data access and computation tasks across many genomic regions before combining the regional results to produce global results.

## RESULTS

### Benchmarking of operating system I/O APIs

Since our work is focused on extracting the maximum I/O performance, we start with benchmarking various operating system I/O APIs. While the POSIX synchronous I/O APIs like *fread()* have been the long-standing standard, newer APIs such as *libaio* and *io_uring* now exist with promises to deliver better performance. Using the *fio* benchmarking utility, we carried out parallel read bandwidth benchmarking on both a Lustre distributed system available locally in our facility, as well as a 4-way NVMe SSD raid0 array available via AWS. We evaluated four APIs on AWS: POSIX synchronous, memory map, *libaio*, and *io_uring*; while dropping *io_uring* on our local Lustre because it is not yet supported by the Linux kernel deployed at our high performance compute cluster. As shown in **Figure 2** (raw data available in **Supplemental Table 1**), the POSIX synchronous APIs consistently outperform other APIs on both Lustre and SSD arrays. Therefore, we chose to use POSIX synchronous APIs for our work.

**Figure 2.**
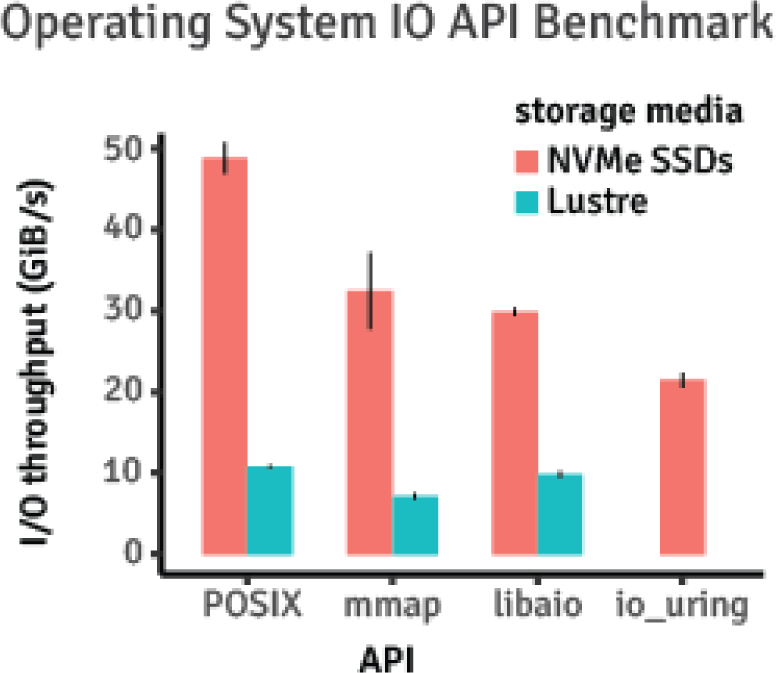
Benchmarking results of various operating system IO APIs. Sequential reads of 4k blocks were carried out using POSIX, memory map (mmap), and libaio on both NVME SSD available on AWS and a Lustre distributed file system available to our local compute cluster. We in addition benchmarked io_uring but only on AWS as it is supported by only recent Linux kernels not yet available at our local facility.

### Performance evaluation strategies

In the next two sections, we describe performance improvements with example algorithms reimplemented using quickBAM. Briefly we describe the benchmarking strategies here with details available in the ***methods*** section. For each algorithm, we benchmarked its performance using two whole genome datasets: Genome In A Bottle(Zook *et al*., 2016) Illumina 2×250 bam files (HG002 and HG004) with a nominal coverage of 75X; and a tumor normal pair 60x Illumina sequencing bam files (Bn2 and Germ1) from a published study(Huang *et al*., 2021). The former dataset is to facilitate result reproduction since it is openly accessible; whereas the latter is to provide a more appropriate tumor context, whose genomes can be highly aberrant. The same tests are carried out separately on Lustre storage at our local cluster and NVMe SSDs on AWS. There are 80 and 96 hyper-threaded cores on our local and AWS servers respectively. Therefore, our Lustre benchmarks contain run configurations of 80, 60, 40, 20, 10, and 1 threads; and NVMe SSD benchmarks contain 96, 72, 48, 24, 12, and 1 threads. For each test, an effective throughput is calculated as total input file size divided by measured total time-till-completion. Each test is repeated 3 times to account for uncontrollable variabilities.

### Proof-of-concept implementation of samtools flagstats and performance evaluation

The first sample program we ported to quickBAM is the utility in samtools called flagstats. Flagstats iterates over the entire BAM file, updating statistics (e.g. number of reads failed QC) with the flags field of each read, and finally printing the statistics when all reads are processed. It is a simple algorithm, however it serves the purpose of demonstrating performance gain via parallelization. Using quickBAM, it is possible to compute separate statistics for each 16 kilo-bases window across the entire genome. This 16k window is chosen because it directly maps to the linear indices in the bam index file. Reads that overlap window boundaries are partitioned into the earlier window to avoid double counting. Since the data structure of flagstats consists of only integer counters, the “gather” stage is a simple summation of these counters from all windows. With a single thread, quickBAM based flagstats and samtools show similar performances (**Figure 3**). However, while samtools benefits little from more than 10 threads, quickBAM allows for a much better scaling. Full timing observations are listed in **Supplemental Table 2**. The quickBAM version of flagstats produces identical results compared to the samtools version. Other algorithms that can potentially be implemented in a similar fashion include, but are not limited to, read counting per fixed genome windows for CNV detection and transcript abundance counting per gene in RNAseq data analysis.

**Figure 3.**
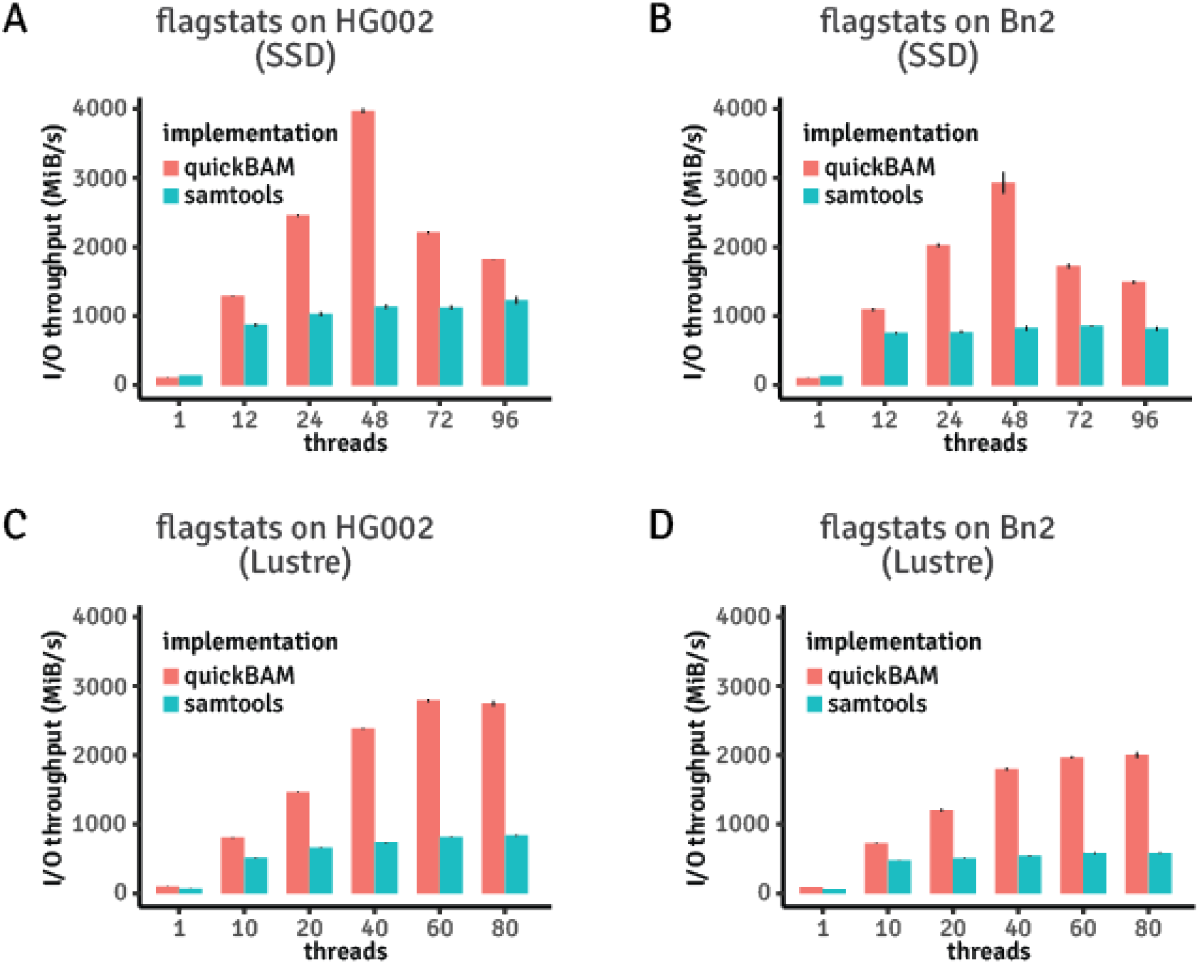
Performance benchmark results of reimplemented flagstats vs stock version. **A)** Results using the GIAB HG002 Illumina 2×250 BAM file with an NVMe SSD array available on AWS. **B)** Results using the rapid autopsy Bn2 sample BAM file with an NVMe SSD array available on AWS. **C)** Results using the GIAB HG002 Illumina 2×250 BAM file with a Lustre distributed file system. **D)** Results using the rapid autopsy Bn2 sample BAM file with a Lustre distributed file system.

### Reimplementation of a real-world, widely used program and performance evaluation

The second sample program we ported to quickBAM is a utility found in the somatic copy number variant detection algorithm FACETS(Shen and Seshan, 2016) called “*snp-pileup*”. *Snp-pileup* takes as input a set of BAM files and a VCF file (commonly the dbSNP published human common polymorphic sites(Sherry *et al*., 1999)), and iterates over positions in the VCF file. At each position, it pulls all the reads from the BAM files overlapping with the position, and extracts the sequencing coverage and variant allele fraction information. Different from the *flagstats* example which parallelizes over multiple, non-overlapping genomic windows, we ported *snp-pileup* to parallelize over groups of consecutive variant positions. As shown in **Figure 4**, quickBAM *snp-pileup* achieved over 1 GiB/s data processing throughput with quickBAM’s built-in multiple input pileup engine, more than 26 times faster than the original implementation which does not support multithreading (HG002 on AWS, 1239.21 MiB/s quickBAM vs 46.38 MiB/s stock). Consequently, using the Genome In A Bottle(Zook *et al*., 2016) HG002 and HG004 Illumina 2×250bp BAM files (242 GiB of data combined), a two samples joint snp-pileup can be finished in 3 minutes 22 seconds (quickBAM), compared to 1 hour 29 minutes (original version). A similar speedup is observed with the tumor normal dataset. Full timing observations are listed in **Supplemental Table 3**. We note that the quickBAM version of *snp-pileup* produces nearly identical results compared to the original version, differing at 217 out of 28.6 million positions (GIAB) and 1879 out of 28.2 million positions (tumor normal). We discuss this in details below. Other algorithms that can potentially be implemented in a similar fashion include single cell sequencing data genotyping and variant calling.

**Figure 4.**
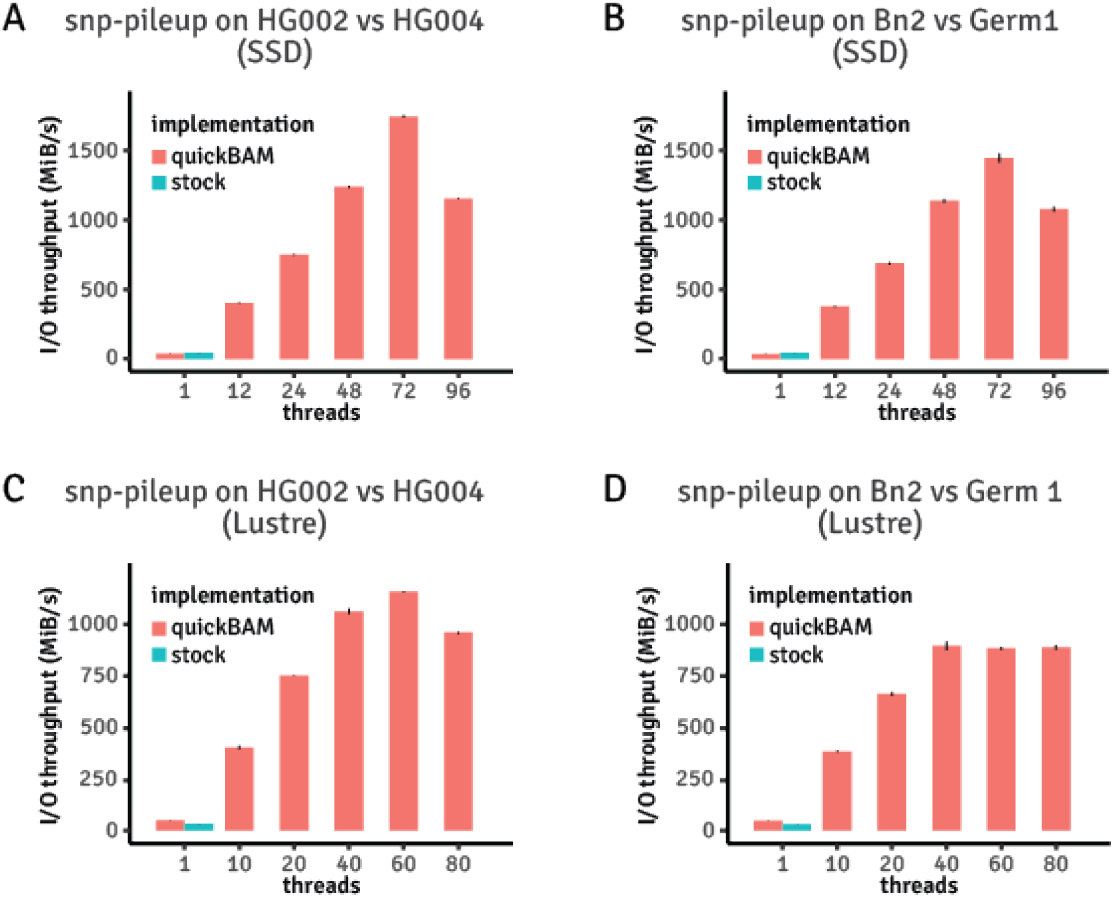
Performance benchmark results of reimplemented snp-pileup vs stock version. Note that the stock implementation of snp-pileup does not support multi-threading. **A)** Results using the GIAB HG002 and HG004 Illumina 2×250 BAM files with an NVMe SSD array available on AWS. **B)** Results using the rapid autopsy Bn2 and Germ1 BAM files with an NVMe SSD array available on AWS. **C)** Results using the GIAB HG002 and HG004 Illumina 2×250 BAM file with a Lustre distributed file system. **D)** Results using the rapid autopsy Bn2 and Germ1 BAM file with a Lustre distributed file system.

We traced the small differences between the original and quickBAM *snp-pileup* to three types of edge cases. The first, which accounts for 10% of the differences, is due to a bug in the stock *snp-pileup* program that would erroneously skip over a location when the input VCF file contains duplicate genomic coordinates, and the first occurrence is not a bi-allelic SNP site. The second, which accounts for 75% of the differences, is due to the difference between how the multiple pileup engines in HTSLIB and quickBAM enforce limits in extremely high coverage regions. HTSLIB uses a “pileup iterator” to which reads are added until the limit is reached, and no more reads will be added until one is removed from the iterator. The pileup engine in quickBAM, however, tries to use every read to update the pileup information at each genomic location. The limit is built into the per-location data structure instead. This generally results in quickBAM counting more even coverage. If there is a demand from the community, we will update the quickBAM multiple pileup engine to behave exactly as HTSLIB. In high coverage regions. For the third type of edge cases (15% of the differences), we verified that quickBAM produced the same results as a third pileup program: *samtools mpileup*. Since the stock *snp-pileup* uses the same HTSLIB multiple pileup engine as *samtools mpileup* does, the exact reason for these differences cannot be determined without putting significant debugging efforts into the original *snp-pileup* program. We thus conclude that these differences are unlikely due to mistakes in quickBAM.

## DISCUSSION

In this manuscript we present quickBAM, a software library for accessing sequence alignments in BAM files with a high degree of parallelization and performance. We achieve this by taking advantage of parallel file access supported by modern storage hardware. As a demonstration of quickBAM’s utility, we ported various types of sequence analysis algorithms, which have shown consistently higher performance than their original implementations. Performance gain is observed with both common on-premise storage solutions such as Lustre, and with new storage hardware such as NVMe SSDs available commonly on the cloud. Porting algorithms to quickBAM offers significant analysis time reduction. As demonstrated by the *snp-pileup* benchmark results, a tumor-normal pair 60X WGS dataset, which took 1.5 hours to process using the original version, can be finished in just under 4 minutes with a quickBAM implementation on the same hardware.

Interestingly, our results show that, while quickBAM allows much higher performance scalability with respect to increasing parallelism, the performance eventually starts to decline as the number of threads approaches the total number of hyperthreaded (with the exception of flagstats on Lustre). This is likely due to oversubscribing system resources, and resulting in software and hardware scheduling overhead exceeding the benefit of increased parallelism. Therefore, it suggests that “sweet-spots” exist for specific hardware / software combinations, and should be determined with trial runs.

Our framework encourages “internal parallelism” i.e. one copy of the analysis program is launched which performs job division and coordinates multi-threading, as opposed to “external parallelism” i.e. many copies of the same analysis programs are launched with each one configured to perform a subset of the total work. There are two benefits of internal parallelism we recognize. First, it is generally easier to program the job division / results gathering processes within the same program space as the actual work routines. We took advantage of this in *snp-pileup* to partition jobs roughly according to the size of data each job spans using the BAM index. And second, since the number of jobs created are independent of the number of threads (with the help of thread pools), it is easier to avoid hardware oversubscription while at the same time benefit from load balancing via job stealing i.e. an idling thread can take jobs from a busy thread to maximize hardware utilization. The trade-off, however, is that internal parallel programs are great at scaling up, but not at scaling out. In future work, we plan to incorporate external parallelism mechanisms such as the OpenMPI software library (Gabriel *et al*., 2004) to make quickBAM even more scalable.

Our work enables many types of sequence analysis software to be accelerated significantly, which in turn benefit time sensitive clinical / research applications such as precision medicine. Our code is open source and publicly available with extensive documentation and sample programs. We plan to actively maintain the project, incorporating further improvements and developing new features according to feedback from the user community.

## ONLINE METHODS

### 1. Core API design principles

We designed our API according to the following principles.

1. The library interface is in C-style data structures and functions that operate on these data structures.

2. To avoid costly abstractions, the data structures are implemented as C structs. The struct members are aligned to the byte arrangement of the records in a BAM or BAI file they represent. The members are intended to be accessed by client programs directly.

3. Functions that perform operations on the data structures are named as <struct_name>_<operation_name> for better code readability.

4. Modern C++11 and C++14 features are used whenever it makes sense. This includes

a. Smart pointers are used for automatic memory management

b. Iterator types in quickBAM often support the ***range-for*** syntax

c. STL algorithms are preferred over explicit loops

d. Auto type deduction is used whenever possible

5. The implementation of quickBAM uses openMP and Intel Thread Building Block (libtbb) for parallelization.

### 2. BAM file partitioning algorithm

QuickBAM speeds up sequence analysis through parallelizing data access and computation over many smaller regions across the genome, which translates to different parts of a BAM file. When the BAM index is available, this is straightforward: spawn a separate task for each 16kb genomic window. This is because BAI directly records the file offset where parsing should start for each 16kb genomic window across the entire genome. Note that 16kb offers the finest level of parallelism. Client code can choose to combine multiple such windows to reduce the number of separate tasks generated. Here we describe in detail when BAI is not available, such as in the case of unsorted BAM files, or the unmapped reads region in an indexed BAM file.

The data stored in a BAM file consists of a series of BGZF compression blocks. A BGZF block contains information about how large the block is. Therefore, it represents a singly linked list data structure i.e. if the location of one BGZF block is already known (e.g. the first one in a BAM file), the location of the next BGZF block can be calculated by *thisBlock*.*offset + thisBlock*.*block_size*. However, when the offset of a BGZF block is not known (e.g. attempting to start reading a BAM file somewhere in the middle), the data structure does not directly indicate where the next BGZF block starts. We use a heuristic algorithm to scan for the next BGZF block using four magic numbers that are guaranteed by the BAM file specification to be present in a BAM file BGZF block: 31(byte 0), 139(byte 1), 66(byte 12), and 67(byte 13). This allows us to scan a BAM file from multiple arbitrary offsets, and find the next valid BGZF blocks. Once a valid BGZF block is located, the rest can be located using *block_size* values.

The bigger challenge of this approach lies in the data stream after decompressing a series of BGZF blocks. That data stream contains the BAM alignment records that do not necessarily align with the beginning of a BGZF block. Since BAM records are similarly a singly linked list, we need to scan the decompressed data to locate the first valid BAM record. This is more difficult because BAM records do not have predictable magic numbers. We instead use properties of a BAM record to heuristically verify if we have found a valid one. We currently implement the following criteria:

- −1 <= refID < n_ref (n_ref is the number of references, part of the BAM header)
- pos == 0 if refID == −1; else pos < l_ref[refID] (l_ref is the lengths of references, part of BAM header)
- −1 <= next_refID < n_ref
- next_pos == 0 if next_refID == −1; else next_pos < l_ref[next_refID]
- read_name[l_read_name − 1] == 0 (this is because read name is NULL terminated)

This set of criteria is derived from the definitions of these properties, and should be agnostic to sequencing technologies / read length / species. Using the datasets mentioned in this manuscript, we have not yet encountered a false positive case. We will optimize our criteria list in future updates if false positive cases are discovered.

With both of these scanning techniques, we are able to parallelize data access across an un-indexed BAM file, or the unmapped reads in an indexed BAM file into which the index does not provide finer partitioning. We note that this approach only applies to read-based algorithms (e.g. flagstats) as position-based algorithms (e.g. snp-pileup) by definition only operates on coordinate-sorted BAM files in mapped regions.

### 3. Benchmark experiments environment

In this section we describe the compute environment setup for our benchmark experiments to facilitate results reproduction.

#### 3.1 Benchmarking using SSD storage array on AWS

We performed benchmark experiments on the Amazon Web Service (AWS) cloud, using a c5d.24xlarge instance and ubuntu 20.04 operating system (AMI ID: ami-03d5c68bab01f3496). This instance type has 96 logic cores, and 192 gigabytes of system memory. Input files are placed on a raid 0 logical volume created from the 4 NVME SSD devices available on this instance, with the following linux commands:

- `mdadm --create /dev/md0 --level=stripe --raid-devices=4 /dev/nvme{1,2,3,4}n1`
- `mkfs -t ext4 /dev/md0`

Because the linux operating system kernel caches all file read operations into unused system memory, the subsequent experiments reading the same files will not incur actual disk operations. To maintain the same conditions between experiments, the kernel caches are manually dropped using the following linux command between benchmark runs

- `echo 3 > /proc/sys/vm/drop_caches`

Note that this is different from disabling cache altogether. We permit caching during a single experiment run so that parallel tasks reading the same sections of a file (though rarely) can benefit from system cache.

Elapsed times are measured using the “gnu time” utility shipped with the operating system.

#### 3.2. Benchmarking using Lustre distributed file storage on local compute clusters

We performed benchmarking experiments using our local high performance compute cluster hosted at the Center for High Performance Computing at the University of Utah. We have access to a 500 terabyte Lustre distributed file system as part of our multi-group shared storage. Lustre is a complex file system that has multiple layers of caching that is beyond the control of an end-user; and it serves multiple research groups and users. Therefore, the exact benchmarking conditions are difficult to reproduce. Our results represent our best effort at minimizing test environment variabilities.

The server that has access to the file system has 80 hyper-threaded cores and 384 gigabytes of system memory. We had exclusive access to this server during our benchmarking experiments. Local cache is cleared between experiments in the same way as described in ***method 3*.*1***.

### 4. Datasets

We used two datasets to conduct all our benchmarking experiments. The first is the publicly and openly available Genome In A Bottle (GIAB)(Zook *et al*., 2016) Ashkenazim Trio Illumina 2×250bp GECh38 BAM files. The inclusion of this dataset is to facilitate result reproduction. The second is a tumor-normal sample pair from a published study(Huang *et al*., 2021), Bn2 and Germ1 specifically, that is under controlled access. While gaining access to this dataset is more tedious, it serves to provide a more appropriate cancer content for tools like FACETS since tumor genomes can be highly aberrant.

### 5. Flagstats benchmark experiment

We ran the flagstats utility as part of samtools v1.16.1 for our benchmark experiments. We used the ‘-@’ parameter to control the number of threads samtools can create.

### 6. Snp-pileup benchmark experiment

We used the VCF recommended by the snp-pileup documentation (ftp://ftp.ncbi.nlm.nih.gov/snp/organisms/human_9606/VCF/00-common_all.vcf.gz) for sites of known polymorphism. For the stock version of snp-pileup, we used the ‘-d 2000’ option to limit the pileup depth, and the ‘-x’ option because the multiple pileup engine in quickBAM does not currently re-adjust base qualities of overlapping mates. We ran the stock snp-pileup with one thread, repeated three times to obtain standard deviations. No further runs were performed since the stock snp-pileup does not support multi-threading.

### DATA AVAILABILITY

No new data were generated or analyzed in support of this research.

GIAB Ashkenazim Trio HG002 and HG004 Illumina 2×250bp novoalign GRCh38 BAM files are available at

- https://ftp-trace.ncbi.nlm.nih.gov/ReferenceSamples/giab/data/AshkenazimTrio/HG002_NA24385_son/NIST_Illumina_2×250bps/novoalign_bams/
- https://ftp-trace.ncbi.nlm.nih.gov/ReferenceSamples/giab/data/AshkenazimTrio/HG004_NA24143_mother/NIST_Illumina_2×250bps/novoalign_bams/

The rapid autopsy tumor normal sample dataset was from a published study(Huang *et al*., 2021). Known polymorphism sites VCF used for the snp-pileup benchmark experiments are available at

- ftp://ftp.ncbi.nlm.nih.gov/snp/organisms/human_9606/VCF/00-common_all.vcf.gz

## Supporting information

Backend API benchmark results

flagstats stock vs quickBAM

snp-pileup stock vs quickBAM

## ACKNOWLEDGEMENTS

A.P., X.H., G.M., and Y.Q. are supported by NIH Grant U24CA209999, and an internal grant from the Center for Genomic Medicine for the development of computational pipelines supporting precision oncology. A.P., G.M., and Y.Q. are supported by NIH Grant 1R21CA271098-01A1.

## AUTHOR CONTRIBUTIONS

Y.Q. conceived the project, contributed to the development of software code, and designed and performed benchmark experiments; and provided support. A.P. contributed to the design, development, and testing of the software code and documentation. X.H. performed data analysis on the benchmarking experiment results. G.M. contributed to the design of the project, and provided support. All authors read and edited the manuscript.

